# Germ cell connectivity enhances cell death in response to DNA damage in *Drosophila* testis

**DOI:** 10.1101/131425

**Authors:** Kevin L. Lu, Yukiko M. Yamashita

## Abstract

Two broadly known characteristics of germ cells in many organisms are their development as a ‘cyst’ of interconnected cells and their high sensitivity to DNA damage. Here we provide evidence that these characteristics are linked, and that interconnectivity is a mechanism that confers to the *Drosophila* testis a high sensitivity to DNA damage. We show that all germ cells within a cyst die simultaneously even when only a subset of them exhibit detectable DNA damage. Compromising connectivity results in cysts in which only a subset of germ cells die upon DNA damage, lowering overall germ cell death. Our data indicate that a death-promoting signal is shared through the intercellular connections of germ cells. Taken together, we propose that intercellular connectivity is a mechanism that uniquely increases the sensitivity of the germline to DNA damage, thereby protecting the integrity of gamete genomes that are passed on to the next generation.

## Introduction

A prevalent feature of germ cell development across species is their proliferation as an interconnected cluster of cells, widely known as a germ cell cyst. In many organisms from insects to humans, germ cells divide with incomplete cytokinesis that results in interconnected cells with shared cytoplasm, leading to cyst formation (Greenbaum, Iwamori, Buchold, & Matzuk, 2011; Haglund, Nezis, & Stenmark, 2011; Pepling, de Cuevas, & Spradling, 1999). During oogenesis, this intercellular connectivity is critical for the process of oocyte specification, allowing only some of the developing germ cells to become oocytes while the others adopt a supportive role (de Cuevas, Lilly, & Spradling, 1997; Lei & Spradling, 2016; Pepling et al., 1999). For example, in the *Drosophila* ovary, four rounds of germ cell divisions with incomplete cytokinesis results in a cyst of 16 interconnected germ cells, where only one becomes an oocyte while the remaining 15 germ cells become nurse cells. During this process, nurse cells support oocyte development by providing their cytoplasmic contents to oocytes via intercellular trafficking (Cox, 2003; de Cuevas et al., 1997; Huynh & St Johnston, 2004). In contrast to oogenesis, where cytoplasmic connectivity has a clear developmental role in oocyte determination, spermatogenesis is a process where all germ cells within a cyst are considered to be equivalent and become mature gametes (Fuller, 1993; Yoshida, 2016). Despite the lack of a ‘nursing mechanism’ during spermatogenesis, intercellular connectivity is widely observed in spermatogenesis from insects to humans (Greenbaum et al., 2011; Yoshida, 2016). While a function for this connectivity has been proposed in post-meiotic spermatids (Braun, Behringer, Peschon, Brinster, & Palmiter, 1989), the biological significance of male germ cell connectivity during pre-meiotic stages of spermatogenesis remains unknown.

Another well-known characteristic of the germline is its extreme sensitivity to DNA damage compared to the soma, with clinical interventions such as radiation or chemotherapy often resulting in impaired fertility (Arnon, Meirow, Lewis-Roness, & Ornoy, 2001; Meistrich, 2013; Oakberg, 1955). It has been postulated that the high sensitivity of the germline to DNA damage is part of a quality control mechanism for the germ cell genome, which is passed on to the next generation (Gunes, Al-Sadaan, & Agarwal, 2015). However, the means by which the germline achieves such a high sensitivity to DNA damage remains unclear.

Here we provide evidence that germ cell connectivity serves as a mechanism to sensitize the germline in response to DNA damage, inducing cell death in the *Drosophila* testis. We show that an entire germ cell cyst undergoes synchronized cell death as a unit even when only a subset of cells within the cyst exhibit detectable DNA damage. Disruption of germ cell connectivity in mutants of the fusome, an organelle that connects the germ cells within a cyst, leads to death of individual germ cells within a cyst in response to DNA damage, reducing overall germ cell death. The sensitivity of a germ cell cyst to DNA damage increases as the number of interconnected germ cells within increases, demonstrating that connectivity serves as a mechanism to confer higher sensitivity to DNA damage. Taken together, we propose that germ cell cyst formation serves as a mechanism to increase the sensitivity of genome surveillance, ensuring the quality of the genome that is passed on to the next generation.

## Results

### Ionizing radiation induces spermatogonial death preferentially at the 16-cell stage

The *Drosophila* testis serves as an excellent model to study germ cell development owing to its well-defined spatiotemporal organization, with spermatogenesis proceeding from the apical tip down the length of the testis. Germline stem cells (GSCs) divide to produce gonialblasts (GBs), which undergo transit-amplifying divisions to become a cyst of 16 interconnected spermatogonia (16-SG) before entering meiosis (Fig. 1A). In our previous study we showed that protein starvation induces SG death, predominantly at the early stages (∼4-SG stage) of SG development (Yang & Yamashita, 2015) (Fig. 1A). Starvation-induced SG death is mediated by apoptosis of somatic cyst cells encapsulating the SGs (Yang & Yamashita, 2015), which breaks the ‘blood-testis-barrier’ and leads to SG death (Fairchild, Smendziuk, & Tanentzapf, 2015; Lim & Fuller, 2012). Though we also noted significant SG death at the 16-SG stage in the course of our previous work, it was independent of nutrient conditions and thus was not the focus of the study (Yang & Yamashita, 2015).

**Figure 1.**
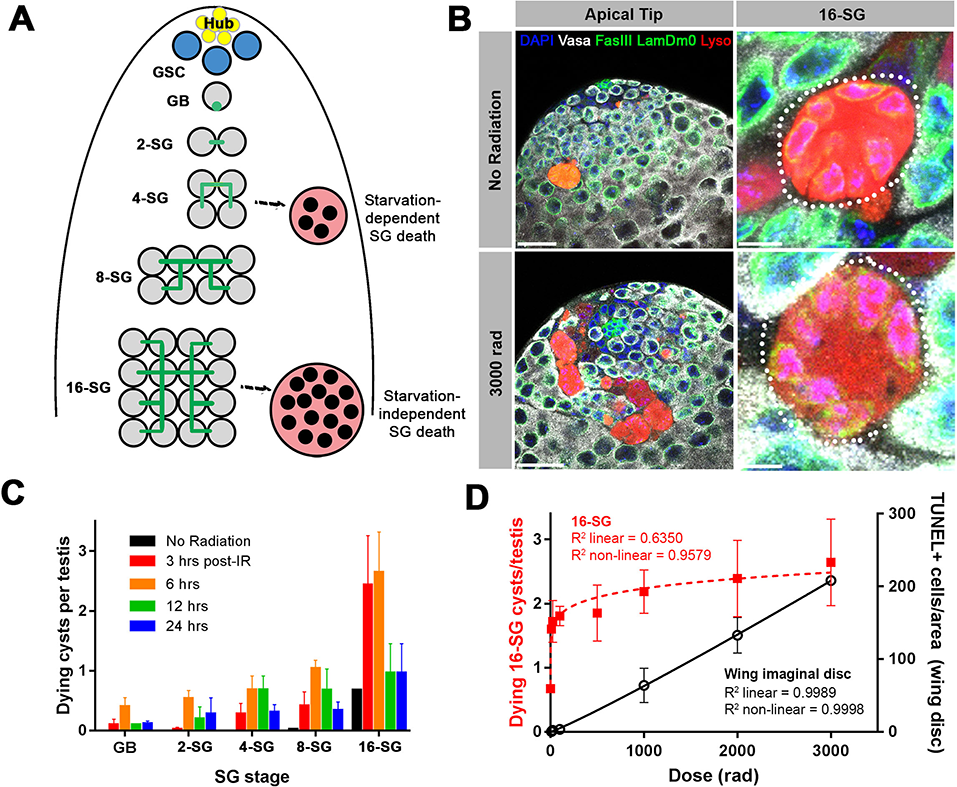
A high level of SG death in response to ionizing radiation. **(A)** Illustration of SG development and germ cell death in the *Drosophila* testis. **(B)** An example of the testis apical tip (left panels) with dying SGs marked by Lysotracker staining in control and irradiated flies. High magnification images of dying 16-SGs (dotted outline) are shown in right panels. Lysotracker (red), Vasa (white), FasIII and Lamin Dm0 (green), and DAPI (blue). Bars: 25μm (left panels), 5 μm (right panels). **(C)** Quantification of dying SG cysts by stage from 3 to 24 hours after 3000 rad (Mean ± SD). Testes sample n ≥ 17, repeated in triplicate. **(D)** Number of Lysotracker-positive 16-SG cysts (red) and TUNEL-positive wing imaginal disc cells (black) 6 hours post-irradiation as a function of radiation dose (Mean ± SD). Testes n ≥ 17 and wing disc n ≥ 3, repeated in triplicate. Best fit lines shown determined by non-linear regression.

In search of the cause of this 16-SG death, we discovered that it can be induced by ionizing radiation (see methods). When adult flies were exposed to ionizing radiation that causes DNA double strand breaks (DSBs), a dramatic induction of SG death was observed (Fig. 1B, C). Dying SGs induced by ionizing radiation were detected by Lysotracker staining, which marks acidified compartments, a hallmark of germ cell death (Yacobi-Sharon, Namdar, & Arama, 2013; Yang & Yamashita, 2015). SG death in control and irradiated flies proceeded in the same manner, where all of the SGs within a cyst die simultaneously by becoming Lysotracker-positive (Fig. 1B). Importantly, in contrast to starvation-induced SG death, which was dependent on somatic cyst cell apoptosis, radiation-induced SG death was not suppressed by inhibiting cyst cell apoptosis (Fig. 1 – Figure supplement 1), suggesting that radiation-induced SG death is a germ cell-intrinsic response.

The frequency of dying SG cysts peaked around 3 to 6 hours after irradiation and decreased by 24 hours post-irradiation (Fig. 1C). Interestingly, we found that ionizing radiation robustly induces cell death at the 16-SG stage, although death of other stages (2-, 4-, 8-cell SGs) was also induced (Fig. 1C and see below). This pattern of SG death held true regardless of the ionizing radiation dose (Fig. 1 -Figure supplement 2). While testing multiple doses of ionizing radiation, we noticed that even exposure to a very low dose of ionizing radiation could dramatically induce death of 16-SGs. By measuring dose-dependent death of 16-SGs at six hours post-irradiation, we found that the 16-SG death induced by increasing radiation was a distinctly non-linear response, quickly reaching a plateau of ∼3 dying 16-SG cysts per testis (Fig. 1D). In comparison, somatic cell death in a defined area of the irradiated wing imaginal disc (Fig. 1 – Figure supplement 3) followed a linear dose-response relationship where an increase in radiation resulted in a proportional increase in cell death (Fig. 1D). These results demonstrate a remarkable sensitivity of 16-SGs to ionizing radiation, compared to somatic cells from imaginal discs.

### All SGs within a cyst die even when only a subset of cells exhibit detectable DNA damage

To gain insight into the cause of the unusual sensitivity of 16-SGs to DNA damage, we evaluated the response of SGs to ionizing radiation at a cell biological level. DSBs result in phosphorylation of the histone H2A variant (γ-H2Av), the *Drosophila* equivalent of mammalian γ-H2AX, reflecting the very early cellular response to DSBs (Madigan, Chotkowski, & Glaser, 2002). Using an anti-γ-H2Av antibody (pS137), we confirmed that γ-H2Av can be robustly detected in SGs following a high dose of ionizing radiation (Fig. 2 – Figure supplement 1). When a low dose of ionizing radiation was used, we frequently observed 16-SG cysts in which only a subset of cells within the cyst possessed detectable γ-H2Av signal but all 16 cells were Lysotracker-positive and dying (Fig. 2A).

**Figure 2.**
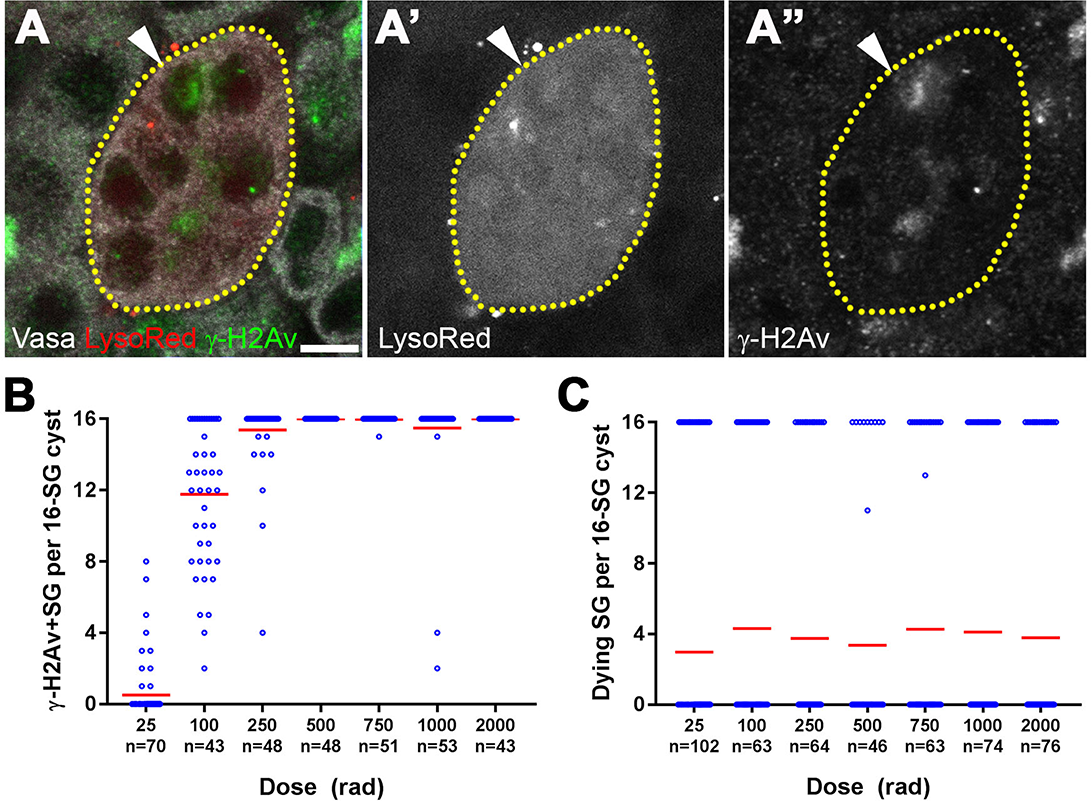
All SGs within a cyst die even when only a fraction of cells exhibit detectable DNA damage. **(A)** An example of a dying 16-SG cyst (yellow dotted outline) with only a subset of SGs containing detectable DNA damage (arrowhead). γ-H2Av (green), Lysotracker (red), Vasa (white). Bar: 5 μm. **(B)** Number of γ-H2Av-positive cells within each 16-SG cyst at various radiation doses. Blue circles, individual data points. Red line, mean. **(C)** Number of Lysotracker-positive cells within each 16-SG cyst. Blue circles, individual data points. Red line, mean.

The fraction of γ-H2Av-positive SGs within each cyst increased gradually with increasing radiation dose irrespective of SG stage (Fig. 2B and Fig. 2 – Figure supplement 2), consistent with the linear nature of ionizing radiation damaging DNA molecules (Ulsh, 2010). However, Lysotracker staining showed that SGs within a cyst were always either all Lysotracker-positive or -negative (Fig. 2C). These results suggest that while DNA damage is induced in individual SGs within a cyst proportional to the dose of radiation, cell death is induced in all of the SGs within the entire cyst, leading to elevated SG death that follows a nonlinear response with increasing dose.

### The fusome is required for synchronized all-or-none SG death within the cysts

The above results led us to hypothesize that all SGs within a cyst might be triggered to die together even when only a subset of SGs within the cyst have detectable DNA damage, explaining the extremely high sensitivity of the germline to DNA damage. In *Drosophila* and other insects, the fusome is a germline-specific membranous organelle that connects the cytoplasm of germ cells within a cyst and mediates intracyst signaling amongst germ cells (Lilly, de Cuevas, & Spradling, 2000; Lin, Yue, & Spradling, 1994). We speculated that if germ cell cysts undergo synchronized cell death by sharing the decision to die, the fusome might mediate the ‘all-or-none’ mode of SG death upon DNA damage.

To examine the role of the fusome in all-or-none SG death upon irradiation, we used RNAi-mediated knockdown of *α-spectrin* and a mutant of *hts*, core components of the fusome (de Cuevas, Lee, & Spradling, 1996; Lilly et al., 2000; Lin et al., 1994) (Fig. 3 – Figure supplement 1). Mutant and control flies were irradiated and their testes were stained with Lysotracker to identify dying SGs in combination with the lipophilic dye FM 4-64 to mark cyst cell membranes, demarcating the boundaries of SG cysts (see methods) (Chiang, Yang, & Yamashita, 2017). In control testes, 16-SG cysts were almost always found to be either completely Lysotracker-positive or -negative under all conditions tested as described above, indicating that 16-SG cysts make an all-or-none death decision (Fig. 3A, B, E, F). Of particular importance, even at a lower dose of radiation (e.g. 100 rad) where only a subset of germ cells exhibit visible γ-H2Av staining (Fig. 2B), the 16-SG cyst was either entirely Lysotracker-negative or -positive. In contrast, *α-spectrin* RNAi and *hts* mutant testes frequently contained 16-SG cysts with a mixture of individual Lysotracker-positive and -negative SGs (Fig. 3C, D, G, H), suggesting that the all-or-none mode of SG death was compromised. Importantly, the mean fraction of 16-SG that died in response to radiation exposure at any dose was significantly reduced when the fusome was disrupted (Fig. 3B, D, F, H, see Supplementary Table S1 for statistics). These data show that the fusome is required for the coordinated death of all SGs within a cyst, and suggests that connectivity of SGs increases overall SG death in response to radiation-induced DNA damage. Moreover, the fact that loss of germ cell connectivity in fusome mutants allows for the survival of some SGs strongly argues against the possibility that SGs without cytologically detectable γ-H2Av are sufficiently damaged to trigger cell death on their own, and that SGs are individually dying. Instead, the death of SGs without detectable γ-H2Av in wild type/control can likely be attributed to a shared signal from other cells, as blockade of intercellular communication in fusome mutants allows for their survival.

**Figure 3.**
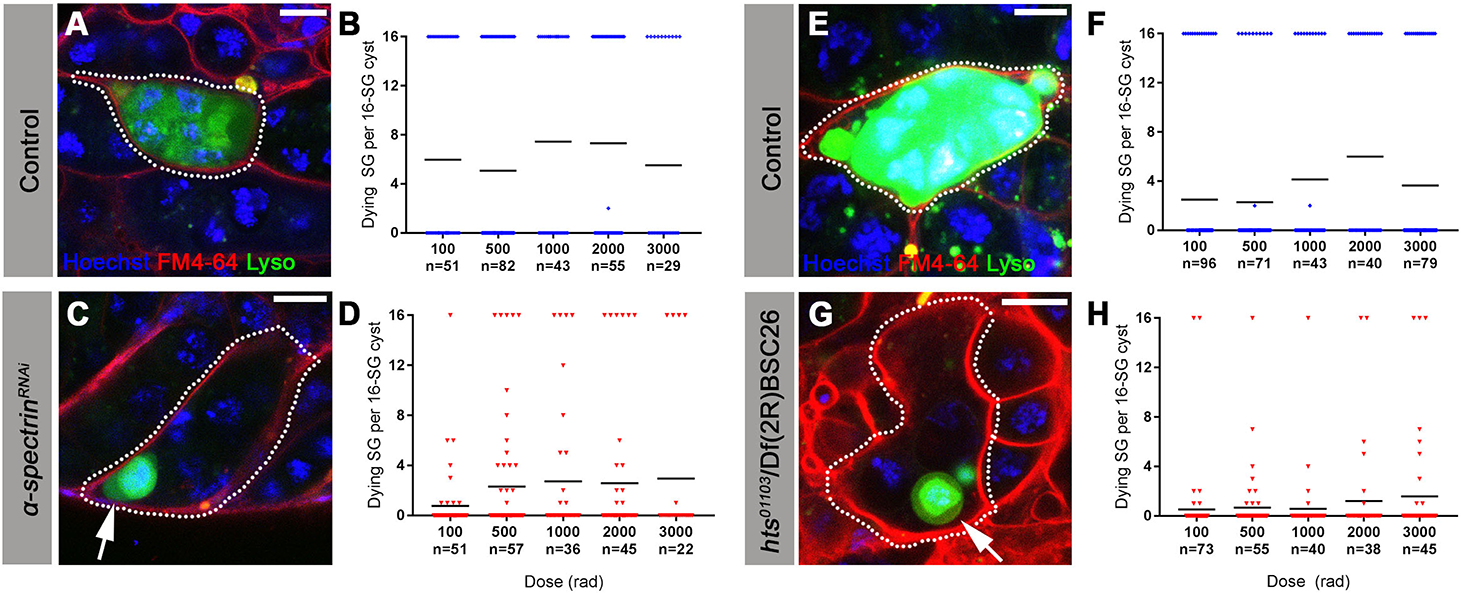
The fusome is required for synchronized all-or-none SG death within a cyst. **(A)** A Lysotracker-positive 16-SG cyst (green, dotted outline) in unfixed control testes, with cyst borders marked by FM 4-64 (red) and SG nuclei marked by Hoechst 33342 (blue). Bar: 7.5 μm. **(B)** Number of Lysotracker-positive cells within each 16-SG cyst of control testes at varying radiation doses. Black line, mean. **(C)** A 16-SG cyst (dotted outline) in *nos-gal4>*UAS*-α-spectrin*^*RNAi*^ testes containing a single Lysotracker-positive SG (arrowhead). Bar: 10 μm. **(D)** Number of Lysotracker-positive cells within each 16-SG cyst of *nos-gal4>*UAS*-α-spectrin*^*RNAi*^ testes at varying radiation doses. Black line, mean. **(E, F)** *hts*^*^01103^*^*/+* control testes. Bar: 5 μm. **(G, H)** *hts*^*^01103^*^/Df(2R)BSC26 mutant testes. Bar: 7.5 μm.

### The mitochondrial protease HtrA2/Omi is required for all-or-none SG death

The above results suggest that intercellular connectivity mediated by the fusome plays a critical role in allowing for all-or-none commitment of SGs to death or survival. Based on these results, we hypothesized that a signal to promote cell death exists that is rapidly transmitted from damaged SGs to others via their intercellular connections.

It has previously been shown that germ cell death in the *Drosophila* testis depends on mitochondria-associated factors rather than effector caspases (Yacobi-Sharon et al., 2013). In response to nuclear DNA damage, the *Drosophila* homolog of the mitochondrial serine protease HtrA2/Omi is cleaved and released from the mitochondrial compartment as part of the mitochondria-associated death pathway, and interacts with components of the apoptotic machinery in the cytoplasm to promote cell death (Igaki et al., 2007; Khan et al., 2008; Tain et al., 2009; Vande Walle, Lamkanfi, & Vandenabeele, 2008).

In *HtrA2/Omi* mutant flies, we frequently observed a mix of Lysotracker-positive and -negative SGs within a single cyst, similar to what was seen in fusome mutants (Fig. 4A). Disruption of the all-or-none mode of SG death within the cyst was observed at any dose of radiation tested (Fig. 4B). Disrupted all-or-none SG death occurred in both heterozygous (*Omi*^*Δ^1^*^/+) and transheterozygous (*Omi*^*Δ^1^*^*/Omi*^*Df*^1^^) conditions, consistent with the previous report that heterozygous conditions exhibit haploinsufficiency in inducing SG death (Yacobi-Sharon et al., 2013). In contrast, wild type control 16-SG cysts maintained their all-or-none mode of SG death (Fig. 4B). Taken together, these data show that HtrA2/Omi is critical for all-or-none SG death within a cyst, and imply a role for HtrA2/Omi in propagating the signal to initiate germ cell death within a SG cyst, possibly through its cycle of cleavage and release from mitochondria.

**Figure 4.**
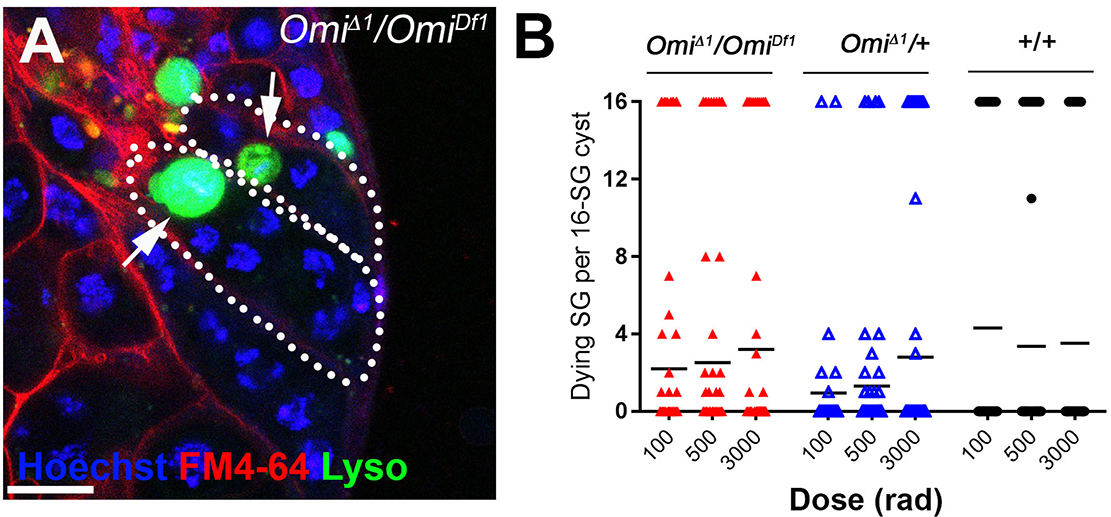
The mitochondrial protease HtrA2/Omi is required for all-or-none SG death. **(A)** SG cysts (dotted outlines) in *Omi*^*Δ^1^*^/*Omi*^*Df*^1^^ mutant testes containing individual Lysotracker-positive SG (arrows). Hoechst 33342 (blue), FM 4-64 (red), Lysotracker (green). Bar: 10 μm. **(B)** Number of Lysotracker-positive SG in each 16-SG cyst in *Omi*^*Δ^1^*^*/Omi*^*Df*^1^^, *Omi*^*Δ^1^*^*/*+, and wild type testes. Black line, mean.

### *p53* and *mnk/chk2* do not regulate the all-or-none mode of SG death

The DNA damage response is a highly conserved pathway controlling cell death and DNA repair (Ciccia & Elledge, 2010; Song, 2005). We thus examined the potential involvement in radiation-induced SG death of the universal DNA damage response pathway components, *mnk/chk2* and *p53,* whose conserved function in DNA damage response in *Drosophila* has been shown (Brodsky et al., 2004; Peters et al., 2002). By using well-characterized loss-of-function alleles (*mnk*^*^6006^*^ and *p53*^*^5^A-^1^-^4^*^) (Takada, Kelkar, & Theurkauf, 2003; Wichmann, Jaklevic, & Su, 2006; Xie & Golic, 2004), we found that these mutants broadly suppress SG death at high and low doses of radiation (Fig. 5D). However, SG death in these mutants maintained an all-or-none pattern (Fig. 5A-C, E). These results suggest that while *p53* and *mnk/chk2* may contribute to SG death via their general role in controlling the DNA damage response as has been described in somatic cells, they do not play a role in mediating the all-or-none pattern of SG death within a cyst that is unique to interconnected germ cells.

**Figure 5.**
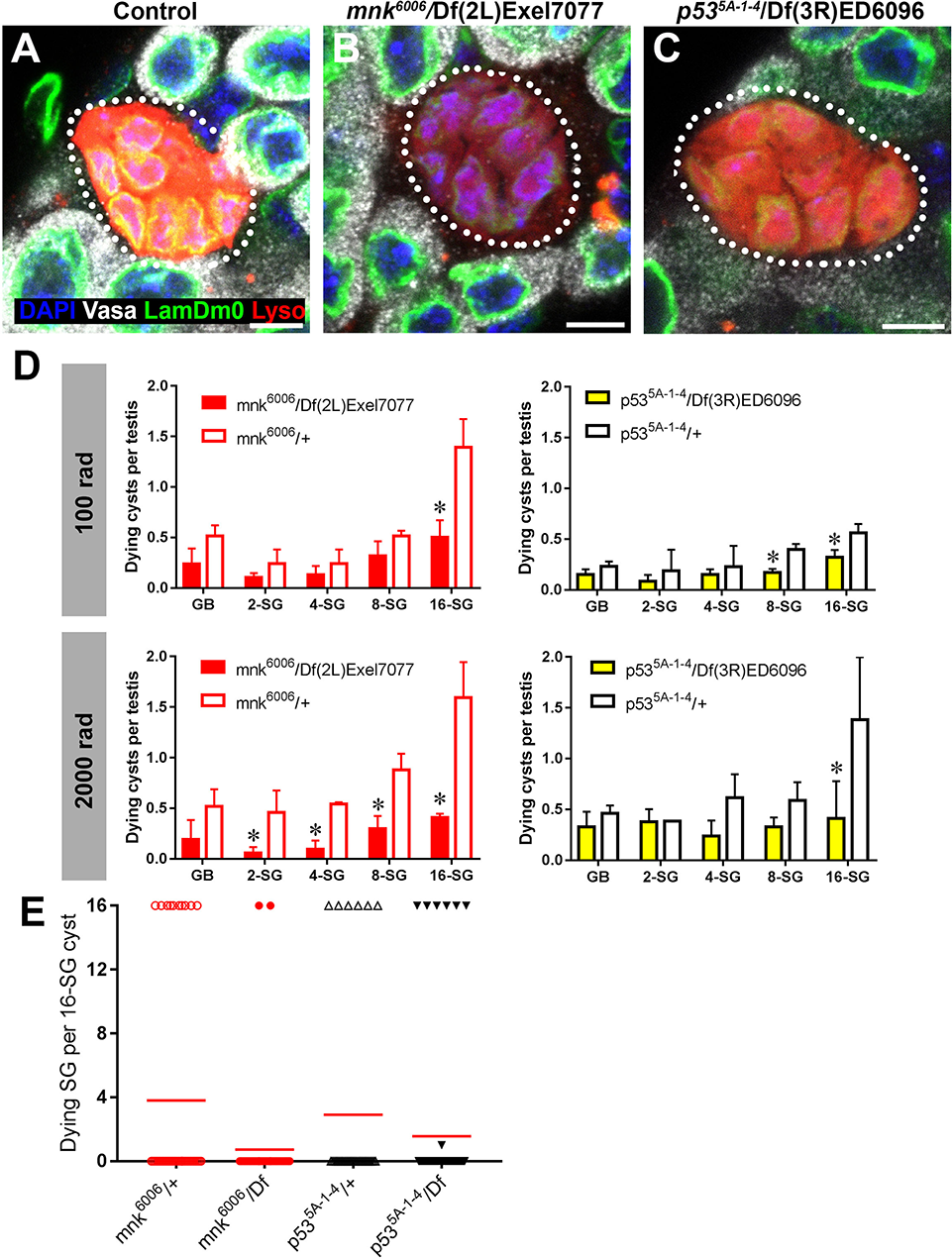
*p53* and *Chk2/mnk* supress SG death but do not regulate the all-or-none mode of SG death. (**A-C**) Examples of dying 16-SGs (dotted outline) in wild-type (A), *mnk*^*^6006^*^*/*Df(2L)Exel7077 mutant (B), and *p53*^*^5^A-^1^-^4^*^/Df(3R)ED6096 mutant (C) testes. Lysotracker (red), Lamin Dm0 (green), DAPI (blue) and Vasa (white). Bars: 5 μm. (**D-G**) SG cyst death by stage in *mnk/chk2* (D, F) and *p53* (E, G) mutants 6 hours after irradiation with 100 rad (D, E) and 2000 rad (F, G) (Mean ± SD, p-value * <0.05 t-test). Fixed, stained samples were used for scoring. Testes sample n ≥ 11 for each genotype, repeated in triplicate. **(H)** Number of Lysotracker-positive SG per 16-SG cyst in *mnk/chk2* and *p53* mutants following 100 rad. Red line, mean. Unfixed samples stained with Lysotracker, FM 4-64, and Hoechst 33342 were used for scoring.

Consistent with the idea that neither *mnk/chk2* or *p53* is responsible for a germline-specific DNA damage response, Mnk/Chk2 or p53 was not upregulated in the germline in response to a low dose of radiation (100 rad), which is normally sufficient to induce robust SG death. Using a polyclonal anti-Mnk/Chk2 antibody (Takada, Collins, & Kurahashi, 2015), we were unable to detect induction of Mnk/Chk2 at a low radiation dose, and even at a high dose Mnk/Chk2 staining was mostly limited to somatic cells (Fig. 5 – Figure supplement 1). Likewise, a p53 transcriptional reporter (Wylie et al., 2014) was not upregulated in response to a low dose of radiation (Fig. 5 – Figure supplement 2). Even at a high dose, the reporter expression was observed only at 24 hours after irradiation, much later than the peak of SG death, which typically happens within a few hours. Collectively, these data indicate that differential expression of p53 or Mnk/Chk2 in SGs is unlikely to account for the high sensitivity of the germline to DNA damage.

### Increasing connectivity of SGs inherently increases sensitivity to DNA damage

The above results suggest that the robust ability of SGs to trigger cell death in response to DNA damage is facilitated by the sharing of signals to die amongst SGs within a cyst, killing SGs that are not sufficiently damaged to commit to cell death on their own. This sharing of death signals is mediated by and dependent on germ cell connectivity. If this is the case, it would be predicted that increasing the connectivity of a SG cyst (the number of interconnected SGs within the cyst) would increase its sensitivity to DNA damage. Indeed, as mentioned above, we observed a trend of 16-SG cysts dying more frequently than 2-, 4-, or 8-SGs (Fig. 1C, Fig 1 – Figure supplement 2). By plotting cell death frequency of all SG stages as a function of increasing radiation dose (Fig. 6A), it becomes clear that the sensitivity of SG cysts correlates with their connectivity, where 8-SGs are less sensitive than 16-SGs but more sensitive than 4-SGs and so on. Interestingly, single-celled GBs, the immediate daughter of GSCs that have not formed any intercellular connections, exhibited an essentially linear increase in death in response to radiation dose, which is reminiscent of somatic imaginal disc cells (Fig. 1D). These results further support the idea that germ cell connectivity plays a key role in increasing the sensitivity of the germline to DNA damage.

**Figure 6.**
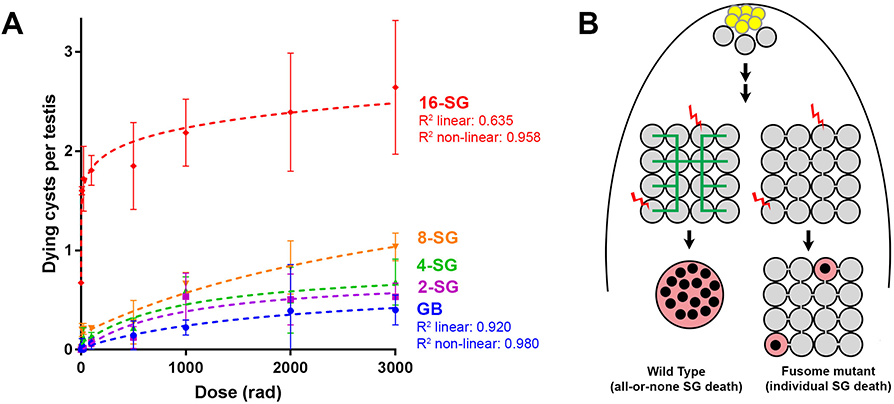
Increasing connectivity confers higher sensitivity to DNA damage. **(A)** Dose-dependent SG death in 2-, 4-, 8-, and 16-SG cysts (Mean ± SD). Best fit lines shown determined by non-linear regression. Testes n ≥ 17, repeated in triplicate. **(B)** Model of SG death enhanced by connectivity.

## Discussion

Our present study may provide a link between two long-standing observations in germ cell biology: 1) the broad conservation of intercellular connectivity (cyst formation) of germ cells and 2) the increased sensitivity of the germline to DNA damage compared to the soma. Our study shows that the connectivity of germ cells is a key mechanism for their ability to robustly induce cell death. We propose that connectivity allows for the sharing of signals that leads to cell death (Fig. 6B). We imagine two possibilities to explain how the signals are shared leading to cell death: when one SG within a cyst decides to die, this ‘decision of death’ might be sent to all the other SGs within the cyst, leading to all-or-none SG death. Alternatively, the signal shared among SGs may be ‘additive’ in nature. In such a scenario, even when none of the individual SGs have sufficient DNA damage to trigger cell death on its own, addition of all damage signaling within the cyst might reach a level sufficient to cause activation of an intracellular cascade that induces cell death. Such a response would effectively lower the threshold of DNA damage needed per cell to trigger germ cell death. Either way, the sharing of signals between SGs through intercellular connections increases their likelihood to die following DNA damage, explaining the unusually high sensitivity of the germline to DNA damage.

According to this model, higher connectivity would confer higher sensitivity to DNA damage: as the connectivity increases, more cells would contribute to detecting any DNA damage the germline may be experiencing. Indeed, our data show a direct correlation between sensitivity to radiation and the increasing connectivity of SG cysts (Fig. 6A). Remarkably, the fact that single-celled, unconnected GBs exhibit an essentially linear death response to increasing radiation suggests that individual germ cells do not have an intrinsically different DNA damage response that accounts for their high sensitivity to DNA damage (Fig. 6A).

A connectivity-based increase in sensitivity to DNA damage also has an important implication in the development of multicellular organisms. To pass on genomes to the next generation, it is critically important for germ cells to have the most stringent mechanisms to prevent deleterious mutations. However, as genome size increases in multicellular organisms, ubiquitously increasing the stringency of genome quality control would result in a high rate of cell death in all tissues, which could compromise the development or survival of organisms. Thus, a multicellular organism would require differential sensitivities to DNA damage between the soma and the germline. The germline would require a more sensitive genome surveillance mechanism to produce gametes with the highest genome quality, whereas the priority of the soma shifts toward survival in order to support development and maintenance of somatic organs. A connectivity-based increase in sensitivity to DNA damage would be a simple method for multicellular organisms to achieve drastically different sensitivities to DNA damage between the soma and germline without having to alter intrinsic damage response pathways. Thus, we speculate that one reason germ cell connectivity has arisen during evolution might be to confer higher sensitivity to DNA damage specifically in the germline.

## Acknowledgements

We thank Drs. Saeko Takada, William Theurkauf, Eli Arama, and John Abrams for reagents. Bloomington Stock Center, the Developmental Studies Hybridoma Bank for reagents, the University of Michigan Experimental Irradiation Core for radiation experiments, Lei Lei, Sue Hammoud and Yamashita lab members for comments on the manuscript.

## Author contributions

K.L. conducted all experiments. K.L. and Y.Y. designed experiments, interpreted results, analyzed the data and wrote the manuscript.

This work was supported the University of Michigan Medical Scientist Training Program, by the University of Michigan Career Training in Reproductive Biology training grant [T32 HD079342] and by NIH Fellowship [F30 AG050398-01] to K.L. The research in the Yamashita laboratory is supported by the Howard Hughes Medical Institute. Y.M.Y. is supported by the MacArthur Foundation. Deposited in PMC for release after 6 months.

## Materials and methods

### Fly husbandry and strains

All fly stocks were raised on standard Bloomington medium at 25°C, and young flies (0 to 2-day old adults) were used for all experiments unless otherwise noted. The following fly stocks were used: *hts*^01103^ (Yuan, Chiang, Cheng, Salzmann, & Yamashita, 2012), *nos-gal4* (Van Doren, Williamson, & Lehmann, 1998), UAS*-α-spectrin*^*RNAi*^ (TRiP.HMC04371), *c587-gal4* (Decotto & Spradling, 2005), UAS-*Diap1* (obtained from the Bloomington Stock Center), *p53*^*^5^A-^1^-^4^*^ (Wichmann et al., 2006; Xie & Golic, 2004), Df(2R)BSC26, Df(3R)ED6096, Df(2L)Exel7077 (obtained from the Bloomington Stock Center), *p53RE-GFP-nls* reporter (Wylie et al., 2014) (a gift of John Abrams, University of Texas Southwestern Medical Center), *Omi*^*Δ^1^*^*, Omi*^*Df*^1^^ (Tain et al., 2009; Yacobi-Sharon et al., 2013) (a gift of Eli Arama, Weizmann Institute of Science)*, mnk*^*^6006^*^ (Takada et al., 2003) (a gift of William Theurkauf, University of Massachusetts Medical School).

### Immunofluorescence staining and microscopy

Immunofluorescence staining of testes was performed as described previously (Cheng et al., 2008). Briefly, testes were dissected in PBS, transferred to 4% formaldehyde in PBS and fixed for 30 minutes. Testes were then washed in PBS-T (PBS containing 0.1% Triton-X) for at least 60 minutes, followed by incubation with primary antibody in 3% bovine serum albumin (BSA) in PBS-T at 4°C overnight. Samples were washed for 60 minutes (three 20-minute washes) in PBS-T, incubated with secondary antibody in 3% BSA in PBS-T at 4**°**C overnight, washed as above, and mounted in VECTASHIELD with DAPI (Vector Labs). The following primary antibodies were used: mouse anti-Adducin-like 1B1 (*hu-li tai shao* – Fly Base) [1:20; Developmental Studies Hybridoma Bank (DSHB); developed by H.D. Lipshitz]; mouse anti-alpha-spectrin 3A9 (1:20; DSHB; developed by R. Dubreuil, T. Byers); rat anti-vasa (1:50; DSHB; developed by A. Spradling), rabbit anti-vasa (1:200; d-26; Santa Cruz Biotechnology), mouse anti-Fasciclin III (1:200; DSHB; developed by C. Goodman), anti-LaminDm0 (1:200; DSHB; developed by P. A. Fisher), rabbit anti-γ-H2AvD pS137 (1:100;Rockland), rabbit anti-mnk (1:100; courtesy of Saeko Takada). Images were taken using a Leica TCS SP8 confocal microscope with 63x oil-immersion objectives (NA=1.4) and processed using Adobe Photoshop software. For detection of germ cell death, testes were stained with Lysotracker Red DND-99 in PBS (1:1000) for 30 minutes prior to formaldehyde fixation. Stages (GB, 2-, 4-, 8-, and 16-SGs) of dying SGs were identified by counting the number of nuclei within the cyst visualized by Lamin Dm0 and DAPI (Chiang et al., 2017). Note that the number of ‘stageable’ dying SGs underrepresents the total population of dying SGs, because nuclear structures disintegrate during later phases of cell death and make it impossible to count the number of SGs within a dying cyst (Chiang et al., 2017). Such ‘unstageable’ SGs were not included in the scoring in this study.

For observation of unfixed samples, testes were dissected directly into PBS and incubated in the dark with the desired dyes for 5 minutes, mounted on slides with PBS and imaged within 10 minutes of dissection. The dyes used in live imaging are: Lysotracker Red DND-99 (1:200) or Lysotracker Green DND-26 (1:200) (Thermo Fisher Scientific), Hoechst 33342 (1:200), and FM4-64FX in PBS (1:200) (Thermo Fisher Scientific). SG stage (2-, 4-, 8-, or 16-SG) was assessed by number of Hoechst-stained nuclei only when using unfixed samples. Note that the scoring of dying SGs is not directly comparable between fixed and unfixed samples (for example, results shown in Fig. 1 vs. Fig. 2), due to the difference in the method of SG staging and timing. In unfixed samples, SG cysts were staged by the number of Hoechst 33342-positive nuclei per cyst marked by FM4-64FX, whereas in fixed samples SG cysts were staged by the number of Lamin Dm0-positive nuclei per cyst.

### Ionizing radiation

For radiation doses of 25-250 rad, a ^137^Cs source was used with a dose rate of approximately 100 rad per minute. Additionally, for radiation doses 100 rad and above, a Philips RT250 model or Kimtron Medical IC-320 orthovoltage unit was used (dose rates of 200 and 400 rad per minute respectively). Dosimetry was carried out using an ionization chamber connected to an electrometer system directly traceable to National Institute of Standards and Technology calibration. The relative biological effectiveness of ^137^Cs and x-ray sources is comparable at lower doses (Fu, Phillips, Heilbron, Ross, & Kane, 1979), and experiments were repeated with both sources for 100-250 rad, which yielded essentially the same results irrespective of radiation source.

### TUNEL Assay

Larval heads with imaginal discs attached were dissected from third instar larvae into PBS, then fixed in 4% formaldehyde in PBS for 30 minutes. Samples were then washed in PBS-T (PBS containing 0.1% Triton-X) for at least 20 minutes, transferred to 100% methanol for 6 minutes with rocking, and washed again for at least 20 minutes in PBS-T. TUNEL assay was then carried out according to manufacturer’s instructions using a Millipore ApopTag Red *In Situ* Apoptosis Detection Kit (S7165). Following washes with PBS-T for 20 minutes, wing imaginal discs were dissected from larval heads and mounted in VECTASHIELD with DAPI (Vector Labs).

### Dose-response best fit regressions

Best fit functions and lines for radiation dose-cell death response curves were generated by using GraphPad Prism 7 and the means-only values at all doses. Non-linear regressions were determined using a four-parameter logistic curve with no constraints on bottom, top, or hillslope and >1500 iterations. Standard linear regression was performed using cell death as a function of radiation dose. Goodness of fit, unadjusted R^2^ value, was determined by 1.0 less the ratio of the regression sum of squares to the total sum of squares, 1 – SS_reg_/SS_tot_.

---

**Fig. 1 – Figure supplement 1.**
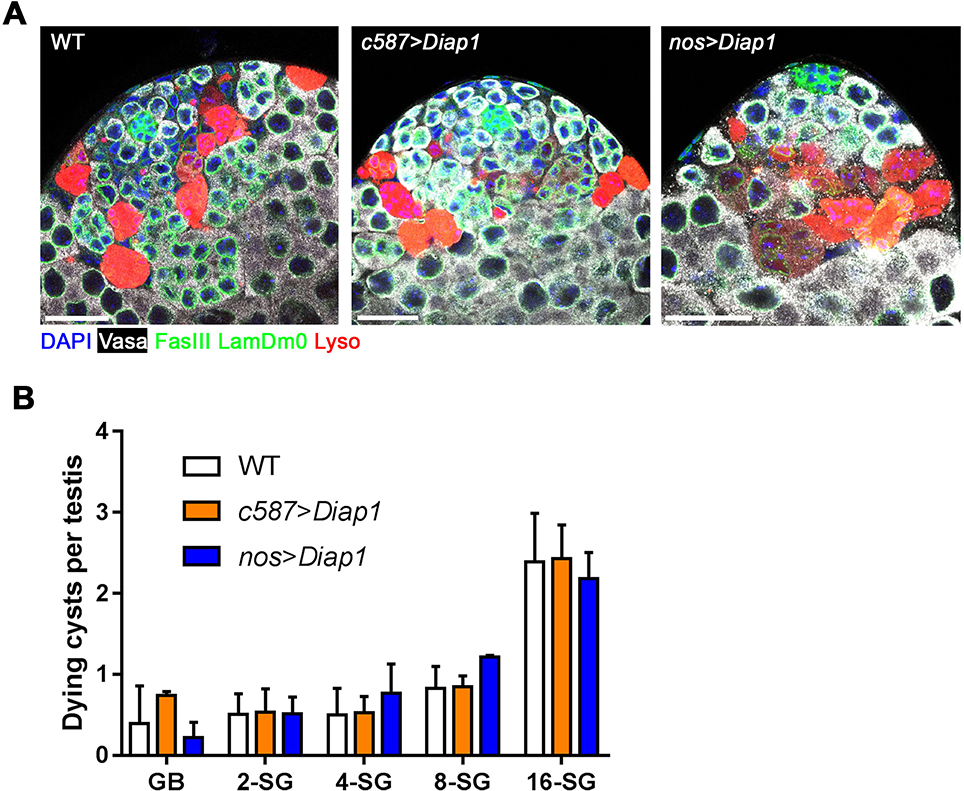
Radiation-induced SG cyst death is independent of somatic cyst cell apoptosis. **(A)** Representative images of testes apical tips from wild type, *c587-gal4>*UAS*-Diap1*, and *nos-gal4>*UAS*-Diap1* expressing flies six hours after 2000 rad. Lysotracker (red), Vasa (white), FasIII and Lamin Dm0 (green), DAPI (blue). Bars: 25 μm. **(B)** Quantification of dying SG cysts by stage from above (Mean ± SD), testes sample n ≥ 15 repeated in triplicate for each. No significant differences at any SG stage between WT vs. *c587*>*Diap1* or *nos*>*Diap1* (p-value * <0.05 t-test).

**Fig. 1 – Figure supplement 2.**
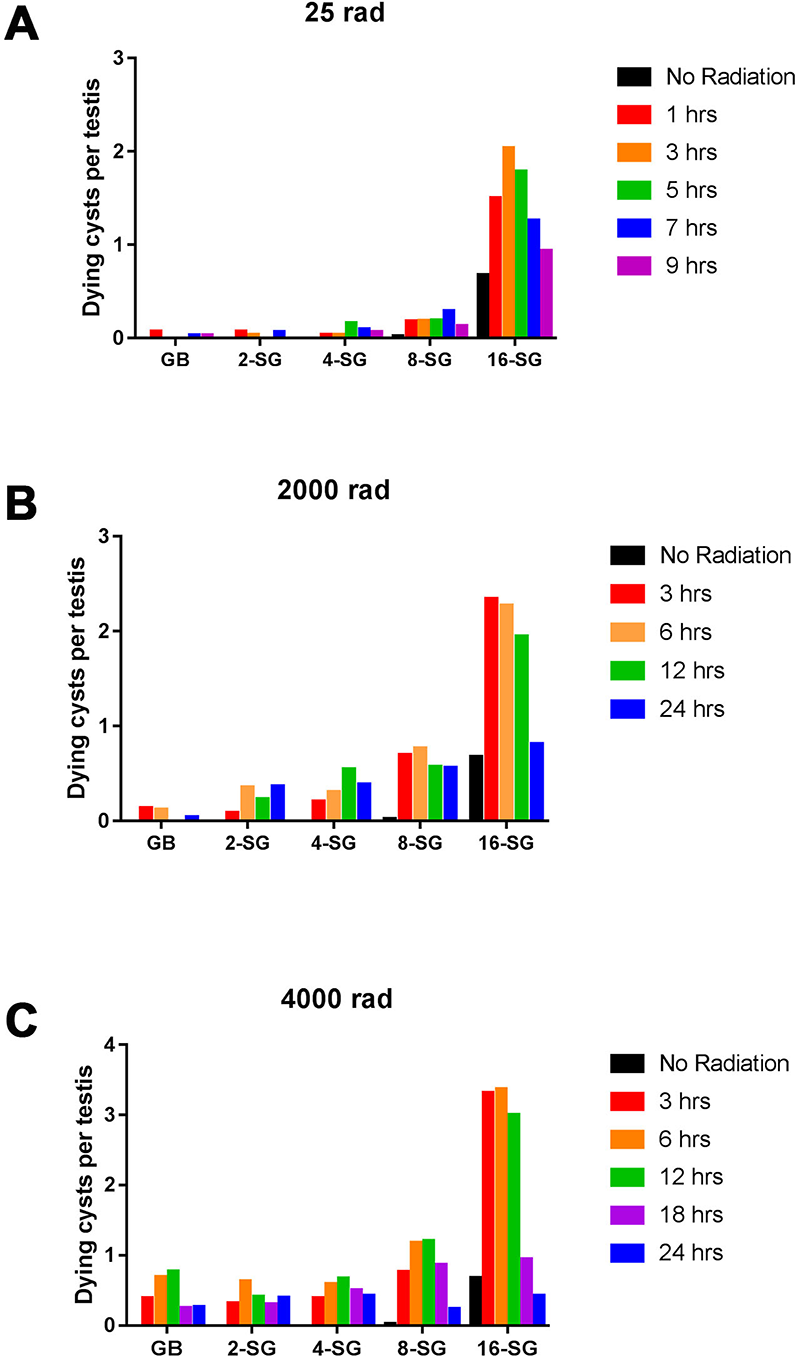
SG death in response to ionizing radiation. Quantification of SG death by stage and time following varying doses of ionizing radiation. **(A)** 25 rad (n = 66), **(B)** 2000 rad (n = 84), **(C)** 4000 rad (n = 68).

**Fig. 1 – Figure supplement 3.**
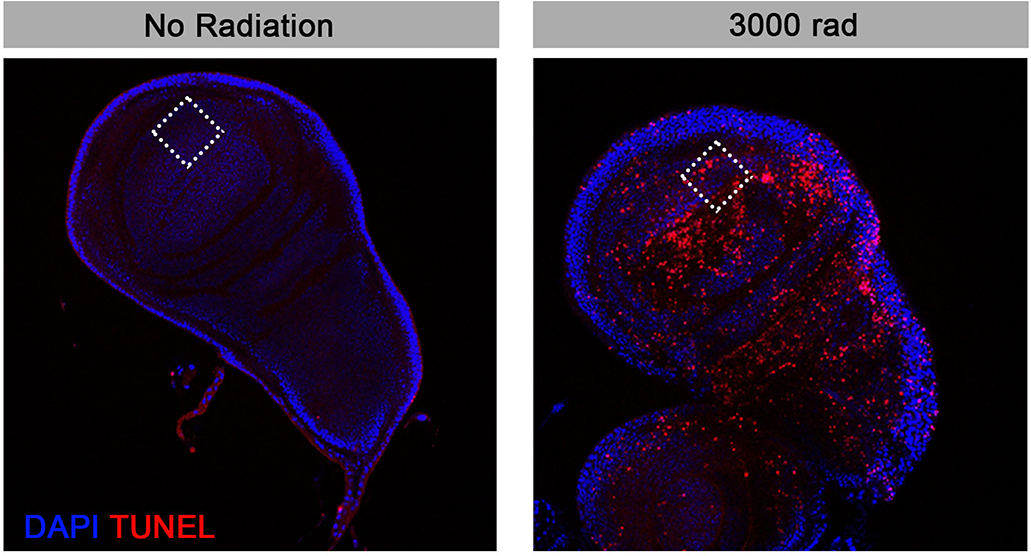
TUNEL staining to detect somatic cell death in response to ionizing radiation. TUNEL-positive foci were quantified in a defined 50 μm^2^ field (dotted square) through entire z-sections in the anterior-ventral compartment of the wing imaginal discs from L3 larvae.

**Fig. 2 – Figure supplement 1.**
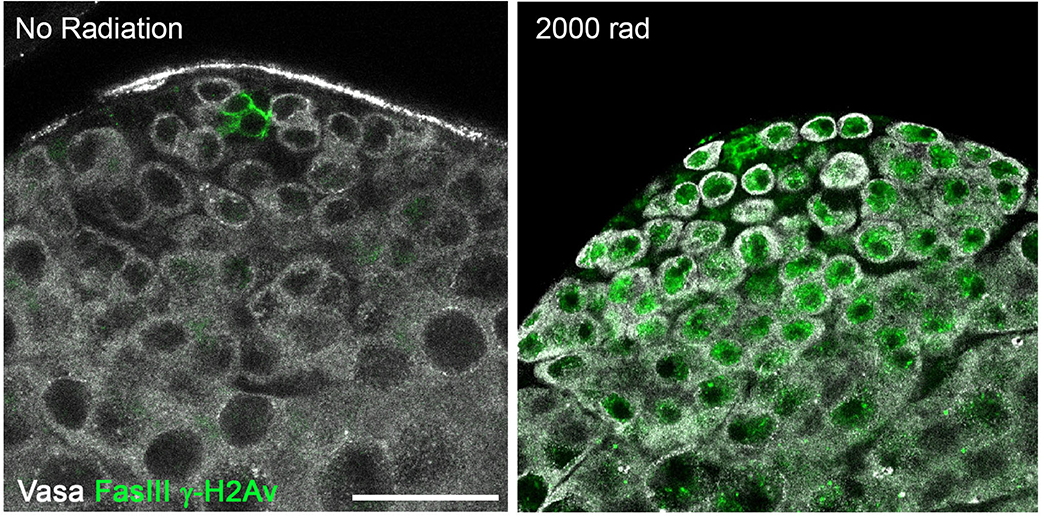
γ-H2Av can be strongly detected in germ cells following irradiation. Representative images of testes apical tips from unirradiated and irradiated flies. Vasa (white), FasIII and γ-H2Av (green). Bar: 25 μm.

**Fig. 2 – Figure supplement 2.**
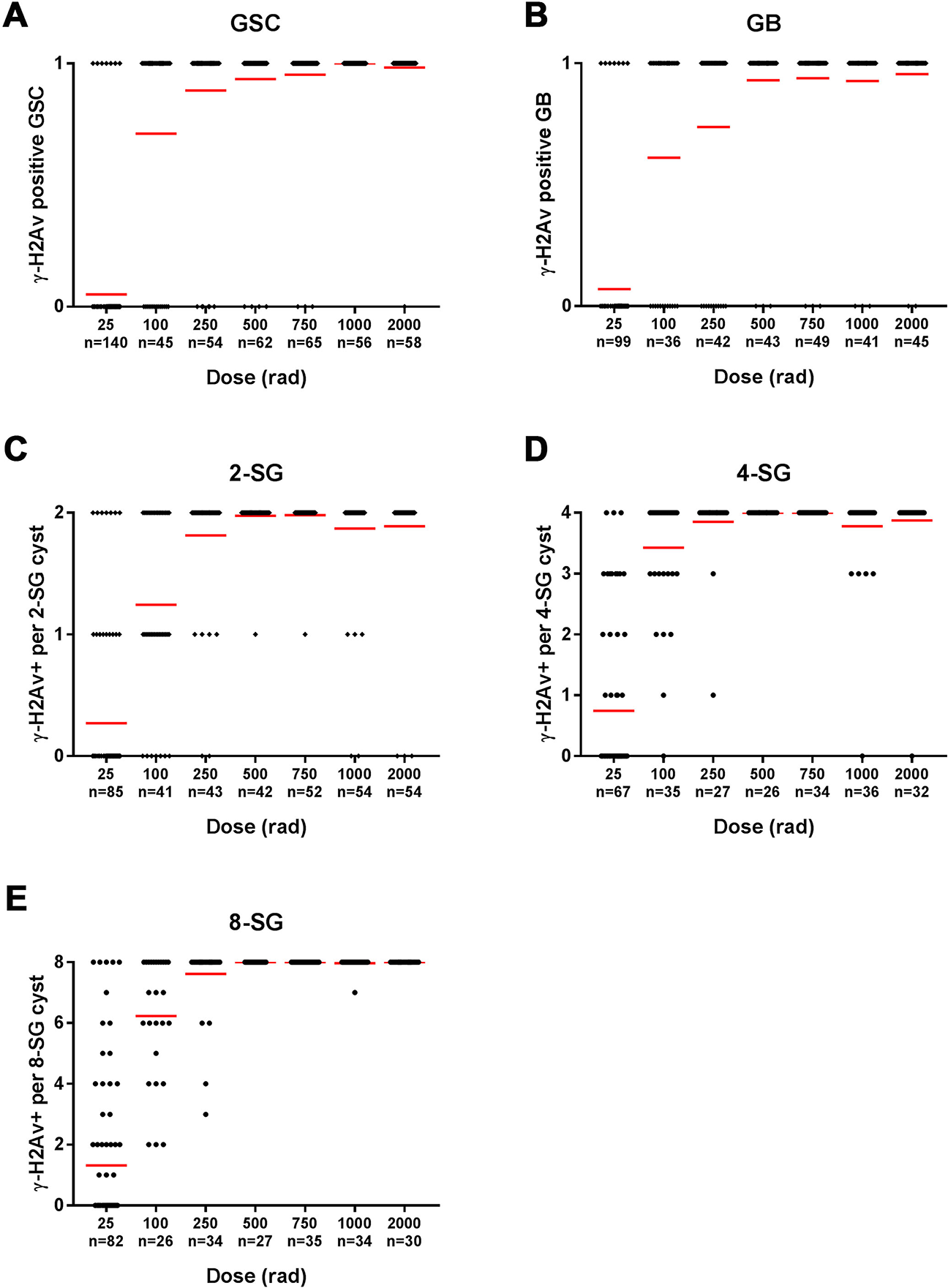
All SG stages show gradual accumulation of γ-H2Av-positive cells with increasing radiation. Number of γ-H2Av-positive cells per cyst at increasing radiation doses in **(A)** germline stem cells **(B)** gonialblasts **(C)** 2-SG **(D)** 4-SG **(E)** 8-SG. Red line, mean.

**Fig. 3 – Figure supplement 1.**
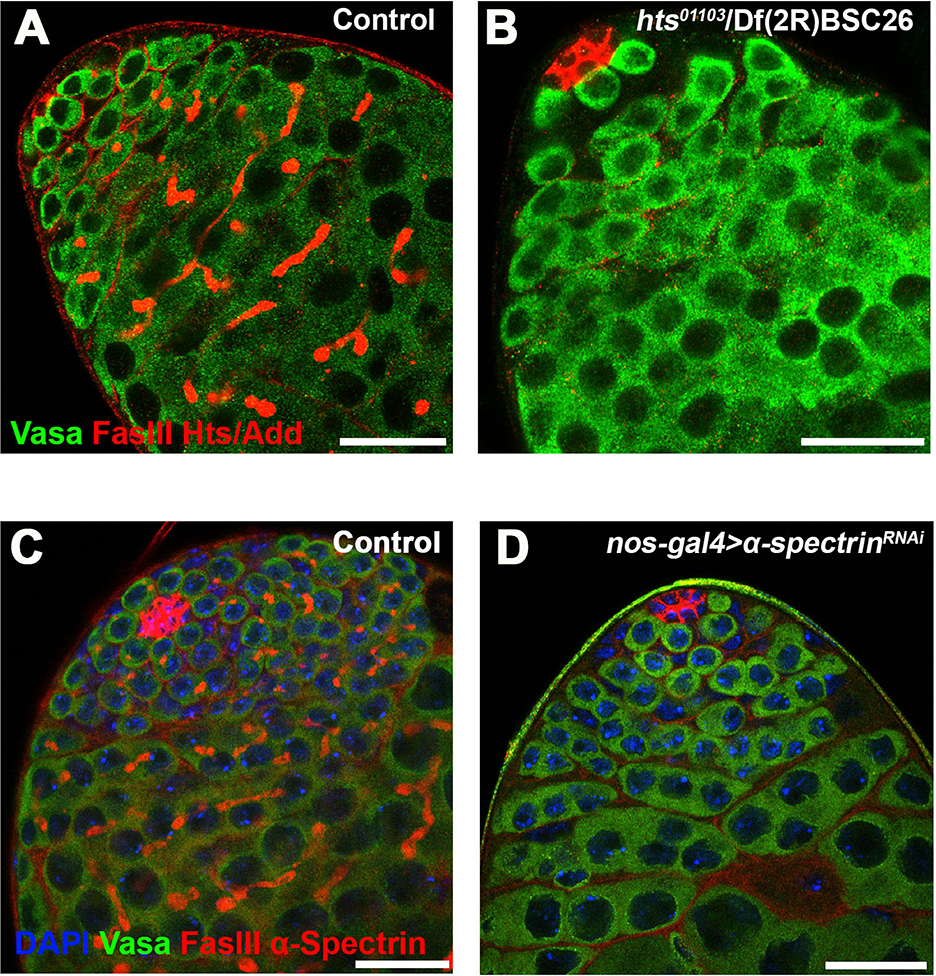
Validation of fusome elimination in *hts* mutant and *α-spectrin* ^*RNAi*^ testes. **(A, B)** Hts/Adducin staining (red) in control (A) and *hts*^*^01103^*^/Df(2R)BSC26 mutant (B) testes. Red: Hts/Add and FasIII. Green: Vasa (germ cells). Bars: 25 μm. Note that *hts* mutation eliminates fusome staining (leaving FasIII staining of the hub cells). **(C, D)** α-Spectrin staining (red) in control (C) and *nos-gal4>UAS-spectrin*^*RNAi*^ (D) testes. α-Spectrin and FasIII (red). Green: Vasa. Blue: DAPI.

**Fig. 5 – Figure supplement 1.**
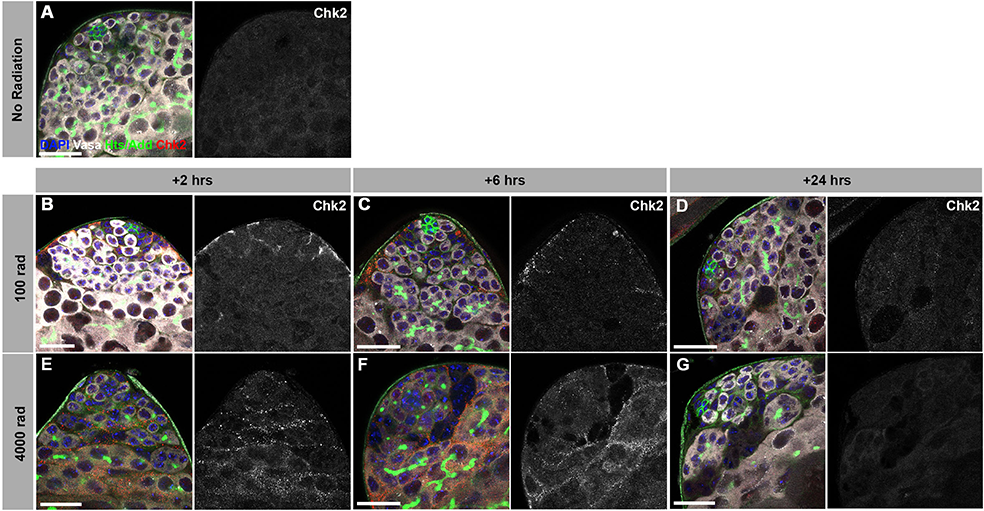
Expression of Mnk/Chk2 in response to ionizing radiation. Testes from unirradiated flies **(A)**, flies irradiated with 100 rad **(B-D)**, or with 4000 rad **(E-G)**, stained for Mnk/Chk2 (red), Vasa (white), Hts (green), and DAPI (blue). Bars: 25 μm. Mnk/Chk2 was detected only after a high dose of radiation, predominantly in the somatic cyst cells surrounding the Vasa-positive germ cells.

**Fig. 5 – Figure supplement 2.**
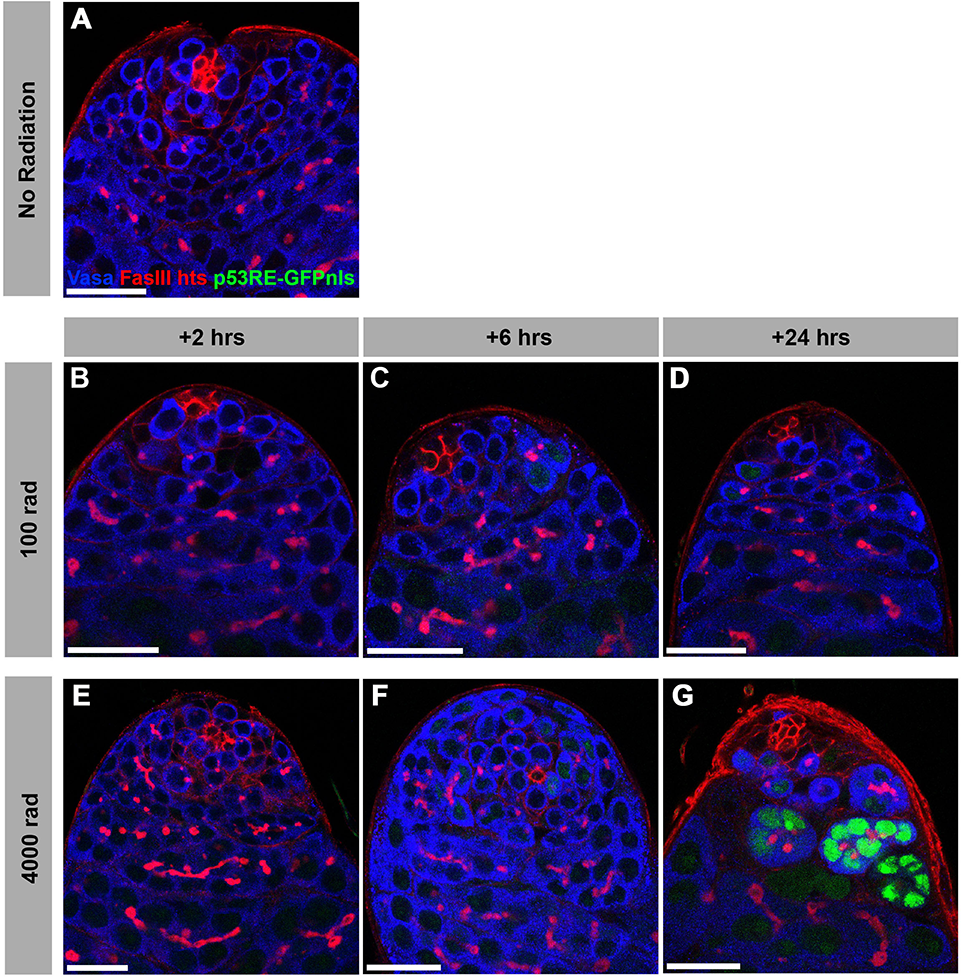
Expression of p53 reporter in response to ionizing radiation. Testes from unirradiated flies **(A)**, flies irradiated with 100 rad **(B-D)**, or with 4000 rad **(E-G)**, stained for p53RE-GFPnls (green), Vasa (blue), Hts/Adducin and FasIII (red). Bars: 25 μm. p53RE-GFPnls was only detectable 24 hours following irradiation at a high dose (4000 rad).

**Supplementary Table S1.** Fraction of 16-SG dying in fusome mutants. Statistics for mean number of dying cells per 16-SG cyst from Fig. 3 in control and fusome mutants. Kolmogrov-Smirnov test for comparing the distribution of two data sets at a given dose of radiation.

| Dose (rad) | Control | 95% CI | *α-Spectrin*<sup>RNAi</sup> | 95% CI | Kolmogorov-Smirnov p-value |
| --- | --- | --- | --- | --- | --- |
| 100 | 5.961 | 3.763<μ<8.158 | 0.7647 | 0.0453<μ<1.484 | 0.0035 |
| 500 | 5.073 | 3.427<μ<6.719 | 2.316 | 1.049<μ<3.583 | 0.0582 |
| 1000 | 7.442 | 4.957<μ<9.927 | 2.722 | 0.8946<μ<4.55 | 0.0147 |
| 2000 | 7.309 | 5.143<μ<9.475 | 2.578 | 0.9354<μ<4.22 | 0.0121 |
| 3000 | 5.517 | 2.573<μ<8.461 | 2.955 | 0.1622<μ<5.747 | 0.8397 |

**Supplementary Table S1. Fraction of 16-SG dying in fusome mutants.**
| Dose (rad) | *hts* Control | 95% CI | *hts* mutant | 95% CI | Kolmogorov-Smirnov p-value |
| --- | --- | --- | --- | --- | --- |
| 100 | 2.5 | 1.317<μ<3.683 | 0.5205 | 0<μ<1.137 | 0.4966 |
| 500 | 2.282 | 0.956<μ<3.607 | 0.6727 | 0.019<μ<1.327 | 0.7395 |
| 1000 | 4.14 | 1.972<μ<6.307 | 0.575 | 0<μ<1.407 | 0.2195 |
| 2000 | 6 | 3.491<μ<8.509 | 1.211 | 0<μ<2.445 | 0.0348 |
| 3000 | 3.646 | 2.133<μ<5.158 | 1.578 | 0.3153<μ<2.84 | 0.4457 |

## References

Arnon, J., Meirow, D., Lewis-Roness, H., & Ornoy, A. (2001). Genetic and teratogenic effects of cancer treatments on gametes and embryos. Human Reproduction Update, 7(4), 394–403. http://doi.org/10.1093/HUMUPD/7.4.394

Braun, R. E., Behringer, R. R., Peschon, J. J., Brinster, R. L., & Palmiter, R. D. (1989). Genetically haploid spermatids are phenotypically diploid. Nature, 337(6205), 373–6. http://doi.org/10.1038/337373a0

Brodsky, M. H., Weinert, B. T., Tsang, G., Rong, Y. S., McGinnis, N. M., Golic, K. G., … Rubin, G. M. (2004). Drosophila melanogaster MNK/Chk2 and p53 regulate multiple DNA repair and apoptotic pathways following DNA damage. Molecular and Cellular Biology, 24(3), 1219–31. Retrieved from http://www.ncbi.nlm.nih.gov/pubmed/14729967

Cheng, J., Türkel, N., Hemati, N., Fuller, M. T., Hunt, A. J., & Yamashita, Y. M. (2008). Centrosome misorientation reduces stem cell division during ageing. Nature, 456(7222), 599–604. http://doi.org/10.1038/nature07386

Chiang, A. C.-Y., Yang, H., & Yamashita, Y. M. (2017). spict, a cyst cell-specific gene, regulates starvation-induced spermatogonial cell death in the Drosophila testis. Scientific Reports, 7, 40245. http://doi.org/10.1038/srep40245

Ciccia, A., & Elledge, S. J. (2010). The DNA Damage Response: Making It Safe to Play with Knives. Molecular Cell, 40(2), 179–204. http://doi.org/10.1016/j.molcel.2010.09.019

Cox, R. T. (2003). A Balbiani body and the fusome mediate mitochondrial inheritance during

Drosophila oogenesis. Development, 130(8), 1579–1590. http://doi.org/10.1242/dev.00365

de Cuevas, M., Lee, J. K., & Spradling, A. C. (1996). alpha-spectrin is required for germline cell division and differentiation in the Drosophila ovary. Development, 122(12).

de Cuevas, M., Lilly, M., & Spradling, A. (1997). GERMLINE CYST FORMATION IN DROSOPHILA. Annual Review of Genetics, 31(1), 405–428. http://doi.org/10.1146/annurev.genet.31.1.405

Decotto, E., & Spradling, A. C. (2005). The Drosophila Ovarian and Testis Stem Cell Niches: Similar Somatic Stem Cells and Signals. Developmental Cell, 9(4), 501–510. http://doi.org/10.1016/j.devcel.2005.08.012

Fairchild, M. J., Smendziuk, C. M., & Tanentzapf, G. (2015). A somatic permeability barrier around the germline is essential for Drosophila spermatogenesis. Development, 142(2), 268–281. http://doi.org/10.1242/dev.114967

Fu, K. K., Phillips, T. L., Heilbron, D. C., Ross, G., & Kane, L. J. (1979). Relative Biological Effectiveness of Low- and High-LET Radiotherapy Beams for Jejunal Crypt Cell Survival at Low Doses Per Fraction. Radiology, 132(1), 205–209. http://doi.org/10.1148/132.1.205

Fuller, M. T. (1993). Spermatogenesis. In The Development of Drosophila melanogaster. In The Development of Drosophila melanogaster (Vol. 1, pp. 71–147). New York: Cold Spring Harbor Laboratory Press.

Greenbaum, M. P., Iwamori, T., Buchold, G. M., & Matzuk, M. M. (2011). Germ cell intercellular bridges. Cold Spring Harbor Perspectives in Biology, 3(8), a005850. http://doi.org/10.1101/cshperspect.a005850

Gunes, S., Al-Sadaan, M., & Agarwal, A. (2015). Spermatogenesis, DNA damage and DNA repair mechanisms in male infertility. Reproductive BioMedicine Online, 31(3), 309–319. http://doi.org/10.1016/j.rbmo.2015.06.010

Haglund, K., Nezis, I. P., & Stenmark, H. (2011). Structure and functions of stable intercellular bridges formed by incomplete cytokinesis during development. Communicative & Integrative Biology, 4(1), 1–9. http://doi.org/10.4161/cib.4.1.13550

Huynh, J.-R., & St Johnston, D. (2004). The Origin of Asymmetry: Early Polarisation of the Drosophila Germline Cyst and Oocyte. Current Biology, 14(11), R438–R449. http://doi.org/10.1016/j.cub.2004.05.040

Igaki, T., Suzuki, Y., Tokushige, N., Aonuma, H., Takahashi, R., & Miura, M. (2007). Evolution of mitochondrial cell death pathway: Proapoptotic role of HtrA2/Omi in Drosophila. Biochemical and Biophysical Research Communications, 356(4), 993–997. http://doi.org/10.1016/j.bbrc.2007.03.079

Khan, F. S., Fujioka, M., Datta, P., Fernandes-Alnemri, T., Jaynes, J. B., & Alnemri, E. S. (2008). The interaction of DIAP1 with dOmi/HtrA2 regulates cell death in Drosophila. Cell Death and Differentiation, 15(6), 1073–83. http://doi.org/10.1038/cdd.2008.19

Lei, L., & Spradling, A. C. (2016). Mouse oocytes differentiate through organelle enrichment from sister cyst germ cells. Science, 352(6281), 95–99. http://doi.org/10.1126/science.aad2156

Lilly, M. A., de Cuevas, M., & Spradling, A. C. (2000). Cyclin A Associates with the Fusome during Germline Cyst Formation in the Drosophila Ovary. Developmental Biology, 218(1), 53–63. http://doi.org/10.1006/dbio.1999.9570

Lim, J. G. Y., & Fuller, M. T. (2012). Somatic cell lineage is required for differentiation and not maintenance of germline stem cells in Drosophila testes. Proceedings of the National Academy of Sciences, 109(45), 18477–18481. http://doi.org/10.1073/pnas.1215516109

Lin, H., Yue, L., & Spradling, A. C. (1994). The Drosophila fusome, a germline-specific organelle, contains membrane skeletal proteins and functions in cyst formation. Development (Cambridge, England), 120(4), |p947–56. Retrieved from http://www.ncbi.nlm.nih.gov/pubmed/7600970

Madigan, J. P., Chotkowski, H. L., & Glaser, R. L. (2002). DNA double-strand break-induced phosphorylation of Drosophila histone variant H2Av helps prevent radiation-induced apoptosis. Nucleic Acids Research, 30(17), 3698–705. Retrieved from http://www.ncbi.nlm.nih.gov/pubmed/12202754

Meistrich, M. L. (2013). Effects of chemotherapy and radiotherapy on spermatogenesis in humans. Fertility and Sterility, 100(5), 1180–1186. http://doi.org/10.1016/j.fertnstert.2013.08.010

Oakberg, E. F. (1955). Sensitivity and Time of Degeneration of Spermatogenic Cells Irradiated in Various Stages of Maturation in the Mouse Sensitivity and Time of Degeneration of Spermatogenic Cells Irradiated in Various Stages of Maturation in the Mouse"’2. Source: Radiation Research, 2(4), 369–391. Retrieved from http://www.jstor.org/stable/3570245

Pepling, M. E., de Cuevas, M., & Spradling, A. C. (1999). Germline cysts: a conserved phase of germ cell development? Trends in Cell Biology, 9(7), 257–62. http://doi.org/10.1016/S0962-8924(99)01594-9

Peters, M., DeLuca, C., Hirao, A., Stambolic, V., Potter, J., Zhou, L., … Mak, T. W. (2002). Chk2 regulates irradiation-induced, p53-mediated apoptosis in Drosophila. Proceedings of the National Academy of Sciences of the United States of America, 99(17), 11305–10. http://doi.org/10.1073/pnas.172382899

Song, Y.-H. (2005). Drosophila melanogaster: a model for the study of DNA damage checkpoint response. Molecules and Cells, 19(2), 167–79. Retrieved from http://www.ncbi.nlm.nih.gov/pubmed/15879698

Tain, L. S., Chowdhury, R. B., Tao, R. N., Plun-Favreau, H., Moisoi, N., Martins, L. M., … Tapon, N. (2009). Drosophila HtrA2 is dispensable for apoptosis but acts downstream of PINK1 independently from Parkin. Cell Death and Differentiation, 16(8), 1118–1125. http://doi.org/10.1038/cdd.2009.23

Takada, S., Collins, E. R., & Kurahashi, K. (2015). The FHA domain determines Drosophila Chk2/Mnk localization to key mitotic structures and is essential for early embryonic DNA damage responses. Molecular Biology of the Cell, 26(10), 1811–28. http://doi.org/10.1091/mbc.E14-07-1238

Takada, S., Kelkar, A., & Theurkauf, W. E. (2003). Drosophila Checkpoint Kinase 2 Couples Centrosome Function and Spindle Assembly to Genomic Integrity. Cell, 113(1), 87–99. http://doi.org/10.1016/S0092-8674(03)00202-2

Ulsh, B. A. (2010). CHECKING THE FOUNDATION: RECENT RADIOBIOLOGY AND THE LINEAR NO-THRESHOLD THEORY. Health Physics, 99(6), 747–758. http://doi.org/10.1097/HP.0b013e3181e32477

Van Doren, M., Williamson, A. L., & Lehmann, R. (1998). Regulation of zygotic gene expression in Drosophila primordial germ cells. Current Biology (Vol. 8). Retrieved from http://www.sciencedirect.com/science/article/pii/S0960982298700910

Vande Walle, L., Lamkanfi, M., & Vandenabeele, P. (2008). The mitochondrial serine protease HtrA2/Omi: an overview. Cell Death and Differentiation, 15(3), 453–460. http://doi.org/10.1038/sj.cdd.4402291

Wichmann, A., Jaklevic, B., & Su, T. T. (2006). Ionizing radiation induces caspase-dependent but Chk2- and p53-independent cell death in Drosophila melanogaster. Proceedings of the National Academy of Sciences of the United States of America, 103(26), 9952–7. http://doi.org/10.1073/pnas.0510528103

Wylie, A., Lu, W.-J., D’Brot, A., Buszczak, M., Abrams, J. M., Abdu, U., … Xu, Y. (2014). p53 activity is selectively licensed in the *Drosophila* stem cell compartment. eLife, 3, e01530.http://doi.org/10.7554/eLife.01530

Xie, H. B., & Golic, K. G. (2004). Gene deletions by ends-in targeting in Drosophila melanogaster. Genetics, 168(3), 1477–89. http://doi.org/10.1534/genetics.104.030882

Yacobi-Sharon, K., Namdar, Y., & Arama, E. (2013). Alternative germ cell death pathway in Drosophila involves HtrA2/Omi, lysosomes, and a caspase-9 counterpart. Developmental Cell, 25(1), 29–42. http://doi.org/10.1016/j.devcel.2013.02.002

Yang, H., & Yamashita, Y. M. (2015). The regulated elimination of transit-amplifying cells preserves tissue homeostasis during protein starvation in Drosophila testis. Development (Cambridge, England), 142(10), 1756–66. http://doi.org/10.1242/dev.122663

Yoshida, S. (2016). From cyst to tubule: innovations in vertebrate spermatogenesis. Wiley Interdisciplinary Reviews: Developmental Biology, 5(1), 119–131. http://doi.org/10.1002/wdev.204

Yuan, H., Chiang, C.-Y. A., Cheng, J., Salzmann, V., & Yamashita, Y. M. (2012). Regulation of cyclin A localization downstream of Par-1 function is critical for the centrosome orientation checkpoint in Drosophila male germline stem cells. Developmental Biology, 361(1), 57–67. http://doi.org/10.1016/j.ydbio.2011.10.010

